# Nanopore amplicon sequencing reveals molecular convergence and local adaptation of opsin genes

**DOI:** 10.1101/2020.06.13.150334

**Authors:** Katherine M. Eaton, Moisés A. Bernal, Nathan J.C. Backenstose, Trevor J. Krabbenhoft

## Abstract

Local adaptation can drive diversification of closely related species across environmental gradients and promote convergence of distantly related taxa that experience similar conditions. We examined a potential case of adaptation to novel visual environments in a species flock (Great Lakes salmonids, genus *Coregonus*) using a new amplicon genotyping protocol on the Oxford Nanopore Flongle. Five visual opsin genes were amplified for individuals of *C. artedi, C. hoyi, C. kiyi*, and *C. zenithicus*. Comparisons revealed species-specific differences in the coding sequence of *rhodopsin* (Tyr261Phe substitution), suggesting local adaptation by *C. kiyi* to the blue-shifted depths of Lake Superior. Parallel evolution and “toggling” at this amino acid residue has occurred several times across the fish tree of life, resulting in identical changes to the visual systems of distantly related taxa across replicated environmental gradients. Our results suggest that ecological differences and local adaptation to distinct visual environments are strong drivers of both evolutionary parallelism and diversification.

## Introduction

Local adaptation to novel environments presents a mechanism that can drive genetic and phenotypic differentiation among closely related organisms. Diversification may occur as populations become locally adapted to distinct conditions, leading to the divergence of traits that are beneficial in each lineage’s preferred environment. Conversely, a trait may be sufficiently advantageous in a particular environment that multiple distantly related taxa converge upon it, in some cases due to the same mutation or amino acid substitution occurring independently, i.e., parallel evolution (Zhang and Kumar 1997, Futuyma and Kirkpatrick 2017). For example, parallel substitutions have occurred in myoglobin in pinnipeds and cetaceans (Romero-Herrera et al. 1978), lysozyme in ruminants and colobine monkeys (Stewart et al. 1987) and rhodopsin in fishes colonizing brackish or freshwater ecosystems (Hill et al. 2019). In this study, we examined a specific case of local adaptation in the teleost visual system that has led to diversification among similar taxa and parallel evolution among distantly related fishes.

Due to their importance in ecological interactions and their dynamic evolutionary history, the evolution of visual pigment genes (i.e. opsins) in marine and freshwater fishes has received considerable attention. The vertebrate visual opsin system is divided into five subgroups – one rod opsin responsible for vision under low light conditions (*rhodopsin*) and four cone opsins responsible for color vision (*long-wave sensitive, short-wave sensitive 1, short-wave sensitive 2*, and *rhodopsin 2*) which are differentiated based on their peak absorbance spectra (Okano et al. 1992). Previous studies have identified two primary mechanisms through which opsin genes can shape the evolution of vision: (1) Single nucleotide polymorphisms (SNPs) causing non-synonymous substitutions in key spectral tuning residues driving adaptation to different light environments (Terai et al. 2002, Marques et al. 2017), and (2) Copy number variation (CNVs) of opsin genes with different spectral tuning (e.g., rod opsin copy number expansions in deep-sea fishes [Musilova et al. 2019], and expansions of cone opsin families in shallow-water fishes [Weadick and Chang 2007]).

The cisco species flock (genus *Coregonus*) of the Laurentian Great Lakes present a well-suited opportunity to study local adaptation of the visual opsin repertoire to novel photic environments based on depth differences (Harrington et al. 2015). The four extant cisco species in Lake Superior show generally low levels of interspecific variation across the genome (Turgeon and Bernatchez 2003, Turgeon et al. 2016, Ackiss et al. 2020) despite considerable differences in depth preferences (Eshenroder et al. 2016, Rosinski et al. 2020). *C. artedi* is typically epilimnetic (10-80 m), *C. hoyi* and *C. zenithicus* are both found at intermediate depths (40-160 m), and *C. kiyi* can be found at depths of 80 to >200 m (Eshenroder et al. 2016). Despite the overall weak genetic divergence among species of the complex, we hypothesized that divergent selection may act to tune opsins to maximally absorb wavelengths of light that penetrate to each species’ preferred depth allowing for prey capture and predator avoidance, leading to measurable genetic differentiation in these species’ opsin genes. Here we assess the evolution of five visual opsins in the *Coregonus* species flock to better understand mechanisms underlying their evolution across a depth gradient. Our aim was to explore both local adaptation to different photic conditions among the closely related *Coregonus* species, and to determine if parallel changes at key spectral tuning sites have occurred among our *Coregonus* species and more distantly related fish taxa.

### New Approaches

Oxford Nanopore sequencing is contributing to a rapidly expanding toolkit for DNA sequencing, owing to low up-front costs, enhanced ability to detect DNA or RNA base modifications, and read lengths limited only by input nucleic acids. Nanopore sequencing allows for straightforward haplotyping, as whole molecules can be sequenced for each amplicon with no need for assembly. This approach has been successfully applied to microbial metabarcoding and pathogen identification (Shin et al. 2016, Moon et al. 2018, Rames and Macdonald 2018), as well as human genotyping (Cornelis et al. 2017, Cornelis et al. 2019). As flow cell quality and base-calling algorithms have improved, the accuracy and functionality of nanopore amplicon sequencing have rapidly expanded. Yet, its application to single nucleotide polymorphism (SNP) genotyping in non-human eukaryotes with large and complex genomes remains relatively unexplored. In particular, a key open question is whether accurate genotypes can be obtained and the coverage depth needed to do so.

In the present study, we sequenced amplicons of five teleost opsin genes in a total of 80 samples on the Oxford Nanopore Flongle device. In combination with the PCR Barcoding Expansion 1-12 (Oxford Nanopore Technologies), we sequenced and genotyped 12 individuals simultaneously on a single Flongle flow cell, following the pipeline shown in Figure 1 (for a complete protocol, see Supplementary File S1). To the best of our knowledge, the present study is one of the first to demonstrate the accuracy and utility of amplicon sequencing with the Oxford Nanopore Flongle for SNP genotyping eukaryotic samples.

**Figure 1.**
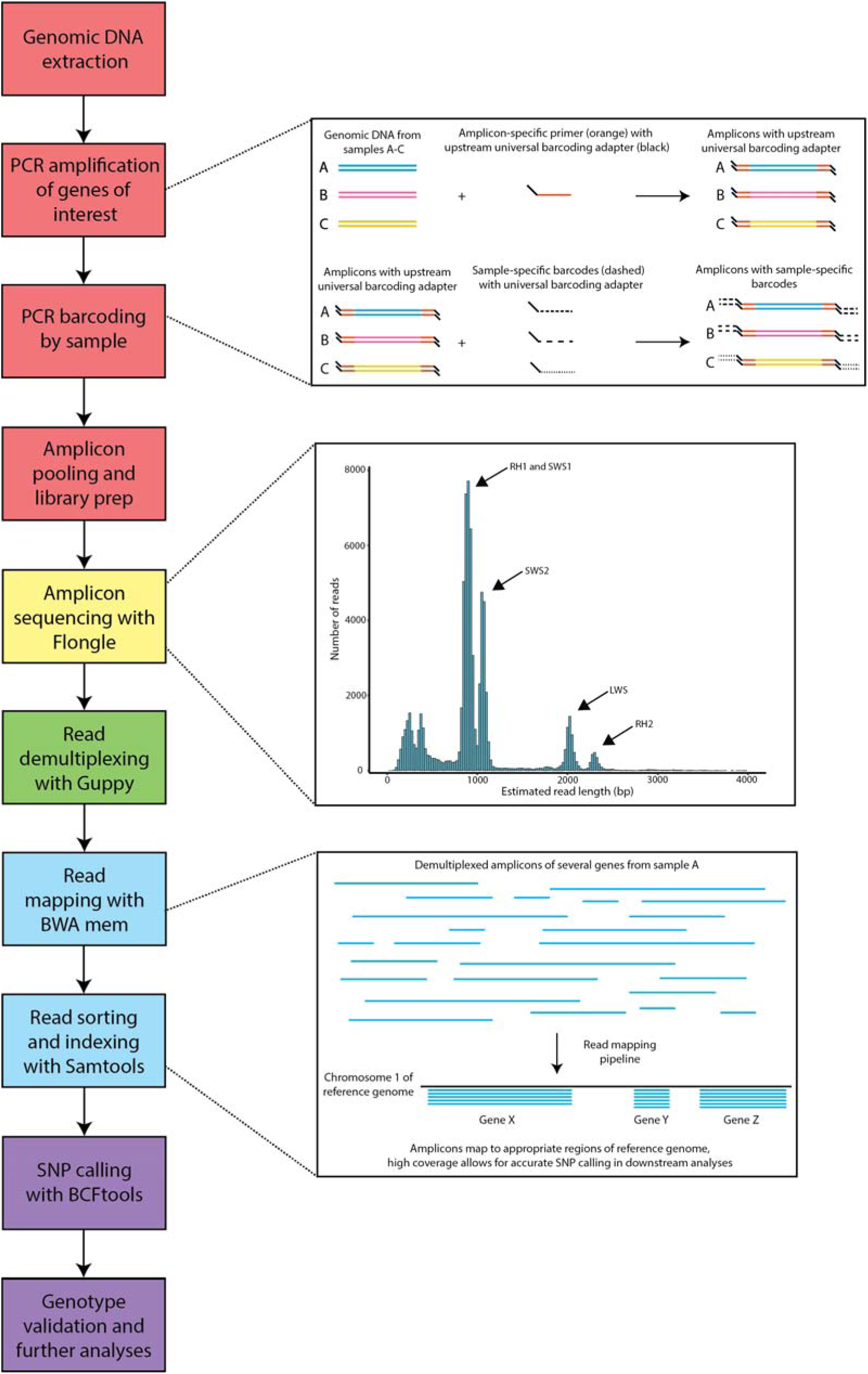
Summary of steps for amplicon sequencing and bioinformatic analyses. Boxes on the left represent individual steps, color-coded based on their phase: red represents sample preparation, yellow represents nanopore sequencing, green represents sample demultiplexing, blue represents read mapping, and purple represents genotyping and analysis. Larger boxes to the right show additional information for each of the steps: the simplified mechanisms by which amplicons are generated and barcoded (top); frequency histogram with read length on the x-axis and number of reads in the y-axis (middle); and how reads are mapped to the reference genome (bottom).

## Results and Discussion

A preliminary assembly of the de novo transcriptome of *Coregonus artedi* (NCBI Bioproject XXXXX) was used as a reference to extract gene sequences of: *long-wave sensitive* (*LWS*), *short-wave sensitive 1* (*SWS1*), *short-wave sensitive 2* (*SWS2*), *rhodopsin* (*RH1*), and *rhodopsin 2* (*RH2*), representing one gene from each teleost opsin subfamily. For each of the five genes of interest, a fragment approximately 700-2100 bp in length was amplified for 18 samples of *C. artedi*, 19 *C. hoyi*, 21 *C. kiyi*, and 16 *C. zenithicus* (Tables 1, S1, S2). All amplicons from a single individual were assigned a specific barcode and were pooled into a library containing genes from 12 samples, which were sequenced simultaneously on a single Flongle flow cell (Figure 1). This process was then repeated until amplicons from all samples were sequenced. After sequencing, sample-specific barcodes were detected and trimmed using Guppy v3.2.4 (Oxford Nanopore Technologies) with the command *guppy_barcoder*, and reads from each sample were mapped with BWA v0.7.17 using the command *bwa mem* (Li 2013), with version one of the *Corgeonus sp. “balchen”* genome assembly as a reference (De-Kayne et al. 2020; GCA_902810595.1). An annotated bash script detailing the entire bioinformatic pipeline is available from Github (https://github.com/KrabbenhoftLab/rhodopsin).

**Table 1.** Average coverage and fragment length for the five amplified opsin genes on the Flongle platform.

| Gene name | Average coverage | Fragment length (bp) |
| --- | --- | --- |
| *LWS* | 722 | 1879 |
| *RH1* | 5409 | 763 |
| *RH2* | 846 | 2079 |
| *SWS1* | 4734 | 815 |
| *SWS2* | 4237 | 960 |

On average, Flongle sequencing runs yielded a total of 206.13 Mb (±166.64 Mb; 26.84-471.50 Mb), with an average of 184,958 reads (±154,877 reads; 23,468-435,138 reads), though yield varied based on flow cell quality (flow cells used were early release and had low starting pore counts). The average sequence N50 was 1,117 bp (±305 bp; 897-1,852 bp), with read length abundances peaking at the approximate lengths of our amplicons (Figure 1). After resequencing genes with low coverage following first-round sequencing, the average coverage was 3,199.58x across all five genes (±4,804.24x, 10.47-31,158.31x; Table 1). Coverage varied slightly by species, but this is likely an artifact of stochastic differences in PCR efficiency and sequencing yield (Table S3). Amplicon reads mapped uniquely (i.e., one genomic region per amplicon) to the *C. sp. “balchen”* genome, providing no evidence for CNVs in opsin genes among *Coregonus* species.

To verify the accuracy of nanopore amplicon genotyping, we performed a rarefaction analysis in which SNPs were called at various levels of coverage (i.e., maximum, 2,000, 1,000, 500, 250, 100, 75, 50, and 25x) in BCFtools v1.9 using the command *bcftools mpileup* (Li 2010, Li 2011, Danecek et al. 2014). The option *-d* was used to specify the maximum per-sample depth. The SNP calls from nanopore data were then compared with Sanger sequences of *rhodopsin* for the same individuals. While accuracy remained high at all sequencing depths (>90%), we found incongruencies in a small proportion of samples between 10x and 75x. Only when reaching 100x coverage were genotypes called with complete accuracy for all individuals, in relation to Sanger sequences. Considering that small errors can impact the results of analyses involving amplicons with few variant sites, we recommend a minimum per-amplicon coverage of 100x for future work.

The genotyping approach used in this study was conservative, as the goal was to assess the coverage needed for accurate genotyping on a Flongle flow cell. Based on our findings, this approach could be used for higher throughput sequencing, which could involve more amplicons, more individuals, or a combination of both. Considering that we generated approximately 200 Mb of sequence data per run, one can calculate the number of individuals and amplicons that can be sequenced simultaneously at 100x using the following formula:

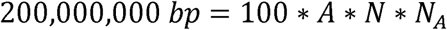

Where *A* is the amplicon size (in bp), *N* is the number of samples to be sequenced simultaneously, and *N*_*A*_ is the number of amplicons to be sequenced per sample. To optimize throughput for the maximum number of samples, the PCR Barcoding Expansion 1-96 (EXP-PBC096, Oxford Nanopore Technologies) can be employed to generate sequence data for 96 samples simultaneously. Assuming an average amplicon size of 1,000 bp, one could sequence 20 amplicons across 96 samples in a single 24-hour Flongle sequencing run for approximately $300, or $0.16 per genotype (Table S4). The use of a MinION flow cell (not analyzed here) would increase output by a factor of ∼16x (based on differences in number of total pores) and reduce the cost per genotype overall. With the growth of nanopore sequencing, these conservative cost estimates are expected to drop in upcoming years.

The average *F*_*ST*_ of SNPs in the five opsins analyzed across four species was 0.055 (Table 2). The only large differences (*F*_*ST*_ > 0.4) were found in four SNPs detected within the coding sequence of *rhodopsin*, with no highly differentiated SNPs among the four cone opsins. This suggests that differences in dim-light vision and changes in *rhodopsin* could be driving local adaptation by depth. Of the four high *F*_*ST*_ SNPs, one (*F*_*ST*_ = 0.44) was synonymous. One SNP (*F*_*ST*_ = 0.44) resulted in a shift from asparagine to histidine at amino acid residue 100, which is located near the C-terminal end of transmembrane helix two, possibly in the extracellular matrix (Figures 2a, 2b; see also Yokoyama 2000). Another (*F*_*ST*_ = 0.44) resulted in a change from valine to isoleucine at residue 255, which is located in transmembrane helix six, facing away from the retinal binding pocket (Figures 2a, 2b, see also Baldwin 1993, Hunt et al. 1996). Neither residue 100 nor 255 are known to be key spectral tuning sites in *rhodopsin* (Yokoyama 2000), but site-directed mutagenesis experiments to determine the effect of these substitutions on the absorbance spectrum should be conducted in the future. All three of these SNPs possess the exact same *F*_*ST*_ and changes in genotype were completely consistent across all samples, suggesting that these sites are tightly linked.

**Table 2.** Pairwise Weir-Cockerham *F*_*ST*_ estimates across species for all genes (above diagonal) and for *rhodopsin* only (below diagonal).

| Species | *C. artedi* | *C. hoyi* | *C. kiyi* | *C. zenithicus* |
| --- | --- | --- | --- | --- |
| *C. artedi* | - | 0.042 | 0.092 | 0.049 |
| *C. hoyi* | 0.26 | - | 0.056 | 0.005 |
| *C. kiyi* | 0.68 | 0.34 | - | 0.068 |
| *C. zenithicus* | 0.26 | 0 | 0.34 | - |

**Figure 2.**
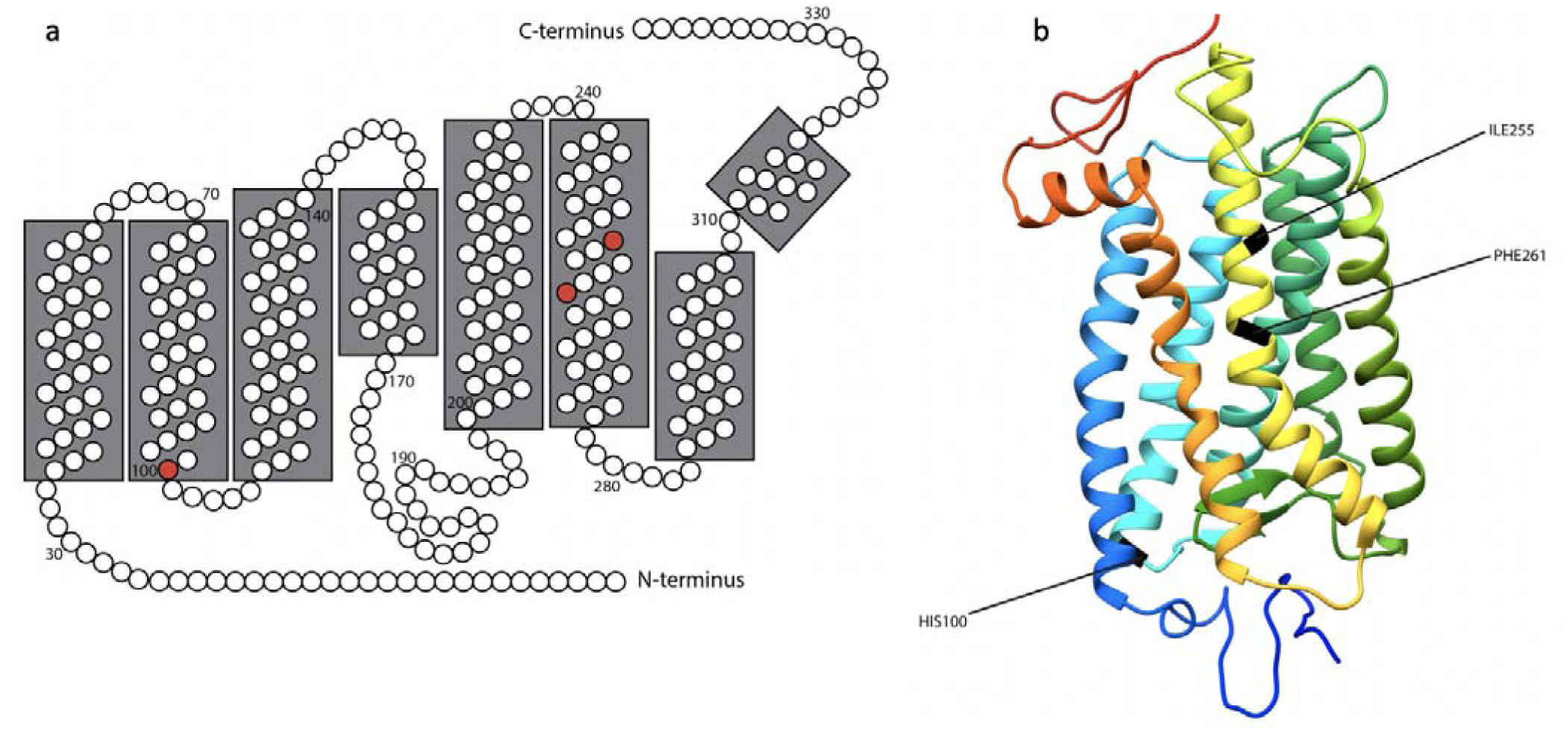
a) 2-D model of *Coregonus artedi* rhodopsin based on a 3-D model generated using PHYRE v2.0 (Kelley et al. 2015) and visualized in UCSF Chimera (Pettersen et al. 2004). Amino acid residues are numbered in order from N-terminus to C-terminus, and amino acid residues 100, 255, and 261 are colored in red. 2-D model was constructed for the specific model obtained for *Coregonus artedi*, following Yokoyama (2000) and Musilova et al. (2019). b) 3-D model of bovine rhodopsin, colored blue (N-terminus) to red (C-terminus). Amino acid residues 100, 255, and 261 have been colored in black, and are labeled accordingly.

The most strongly segregating SNP (*F*_*ST*_ = 0.88) occurred at amino acid residue 261 of *rhodopsin*, which is located in transmembrane helix six, facing the retinal binding pocket of the protein (Figures 2a, 2b, see also Baldwin 1993, Hunt et al. 1996, Yokoyama 2000). *Coregonus artedi, C. hoyi*, and *C. zenithicus*, inhabitants of a red-shifted light environment, were primarily homozygous for tyrosine (Figure 3). This amino acid substitution is known to cause an 8 nm red-shift in the absorbance spectrum (Yokoyama et al. 1995). Meanwhile, *C. kiyi*, which inhabits the blue-shifted deeper waters of Lake Superior, was completely homozygous for phenylalanine, which does not produce a similar red-shift in photic absorbance (Figure 3; Yokoyama et al. 1995). Genotypic associations at this locus vary consistently across the depth gradient (Figure 3), providing evidence that *C. kiyi* is adapted to life in deep water after evolving from shallow-water ancestors. This hypothesis is further corroborated by phenotypic data, as *C. kiyi* have significantly larger eye diameters (as a proportion of total head length) than *C. artedi* (p < 0.001), *C. zenithicus* (p < 0.001), and *C. hoyi* (p < 0.001), consistent with Eshenroder et al. (2016) (Figure S1). The predictable variation of both genetic and morphological traits along the axis of the depth gradient provides key evidence that local adaptation by depth accompanies diversification of Lake Superior ciscoes (Figure S2).

**Figure 3.**
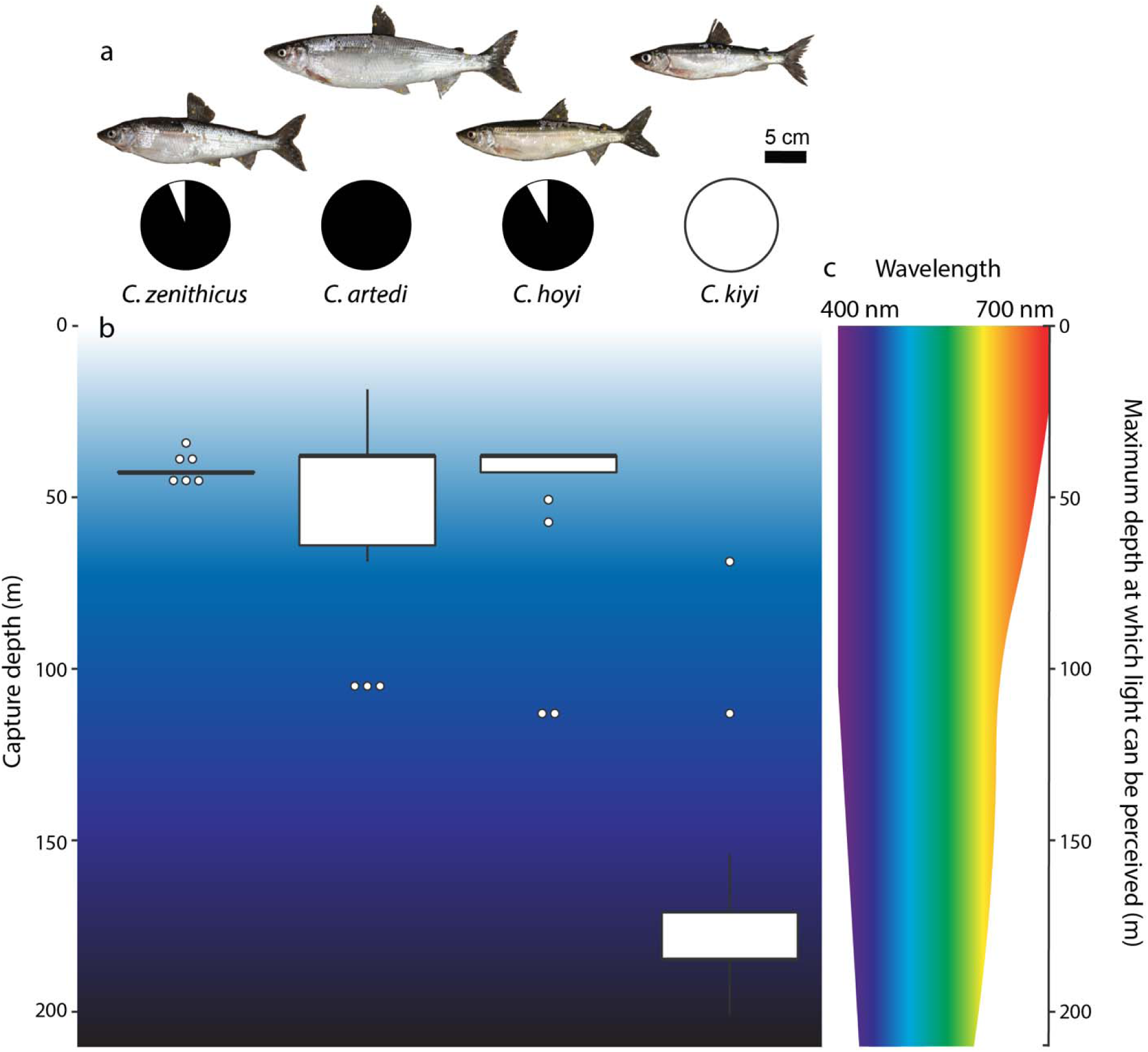
a) The four cisco species included in this study: *Coregonus zenithicus, C. artedi, C. hoyi*, and *C. kiyi* (left → right). Pie charts below each photo indicate the allele frequency at residue 261 of *rhodopsin*, where black represents the allele coding for tyrosine and white represents the allele coding for phenylalanine. b) Boxplots indicating the approximate capture depths of samples from each of the four species. c) Visible light spectrum, from approximately 400-700 nm wavelength. The narrowing of the spectrum with increased depth shows how the ability of organisms to perceive certain wavelengths of light diminishes with increasing depth, particularly with red and orange light (following Harrington et al. 2015).

Hill et al. (2019) examined the shift between the two aforementioned amino acids at *rhodopsin* residue 261 in a deep phylogenetic context, suggesting that many lineages, including salmonids, are likely derived from a marine ancestor possessing the allele encoding the blue-shift-associated 261Phe. Additionally, Hill et al. (2019) found that fish lineages which have undergone a habitat change from blue-shifted marine waters to red-shifted brackish or freshwater have independently converged on the red-shift-associated 261Tyr phenotype over 20 times across the fish tree of life. Here, we show that the exact same substitution has occurred in Great Lakes ciscoes, as the 261Tyr phenotype is predominant among *Coregonus artedi, C. hoyi*, and *C. zenithicus*, which inhabit the red-shifted shallow water of Lake Superior. This finding supports the hypothesis that local adaptation to a novel visual environment is driving parallel molecular evolution across the fish tree of life. Interestingly, it appears that deep-water *C. kiyi* has undergone a reversal to the blue-shifted marine ancestral state (261Phe) after more than 100 million years indicating that *rhodopsin* residue 261 may be able to “toggle” (*sensu* Delport et al. [2008]) between these two amino acids depending on what is advantageous in a particular photic environment, even across incredibly long time scales.

## Conclusions

The present study provides evidence of the utility of the Oxford Nanopore Flongle device for genotyping complex eukaryotic samples by long-read amplicon sequencing. The protocol described is simple and reliable, and offers the promise of rapid, low-cost genotyping in non-model organisms. This methodology was employed here to understand the genetic basis of local adaptation and ecological differentiation among Great Lakes ciscoes.

The results of this study indicate that local adaptation to distinct visual environments is associated with genetic and morphological differentiation among the closely related ciscoes of the Great Lakes. The identification of several high *F*_*ST*_ SNPs in *rhodopsin*, including Phe261Tyr, is particularly relevant, as the shifts between these two amino acids at residue 261 are identical to those observed across similar depth gradients in phylogenetically-distant fishes (Hill et al. 2019). This result suggests that evolutionary parallelism via single-nucleotide changes at this site is driving phenotypic covergence of distantly related groups exposed to similar photic environments. Additionally, the discovery of a reversal to the ancient ancestral state in *C. kiyi* at this site provides evidence of genetic toggling, whereby organisms may be able to transition bi-directionally between different states at this site in response to environmental pressures. This result is striking because the genetic background is persumably very different across these taxa after more than 100 million years of divergence. In addition, the potential for epistatic interactions is expected to be increased across such deep phylogenetic splits, further reducing the likelihood of parallel evolution (Storz 2016). The observation of amino acid toggling in Great Lakes *Coregonus* species stands in stark contrast to the general prediction that evolution can only reverse itself after short time periods (Storz 2016; Blount et al. 2018).

## Acknowledgements

We thank the U.S. Geological Survey Research Vessel Kiyi Captain Joe Walters, First Mate Keith Peterson, and Engineer Charles Carrier, as well as Dan Yule, Mark Vinson, Lori Evrard, Anders Nyman, and the USGS Lake Superior Biological Station and Ontario Ministry of Natural Resources and Forestry for assistance with sample collection. Jessie Pelosi provided critical assistance with data analysis. Mary Alice Coffroth, Tianying Lan, Wendylee Stott, Andrew Muir, Thomas Dowling, Hannah Waterman, Victor Albert, Vincent Lynch, and Omer Gokcumen provided valuable assistance or feedback on the study. Jessica Poulin coordinated the University at Buffalo Honors Program in Biological Sciences that facilitated this research. This work was supported by the Great Lakes Fishery Commission (Award #2018_KRA_44073 to TJK), the University at Buffalo Department of Biological Sciences (Philip G. Miles Fellowship to KME), and the University at Buffalo Honors College (Award to KME).

