## Supplemental Document for "Nanopore amplicon sequencing reveals molecular convergence and local adaptation of opsin genes"

**Supplementary Materials and Methods:**

*Sampling*

Samples were collected from Lake Superior in 2015 and 2019 (Table S1). Upon sampling, fish were assigned to species based on morphological characteristics, following Eshenroder (2016), yielding 18 *Coregonus artedi*, 19 *C. hoyi*, 21 *C. kiyi*, and 16 *C. zenithicus*. Samples obtained in 2015 were dissected, and gill tissue was preserved in RNALater and subsequently stored at -20°C until ready for future analyses. Fishes from 2019 were frozen whole and stored at -80°C.

*Candidate Gene Identification*

Based on the known differences in depth preference among the four studied *Coregonus* species, we hypothesized functional variation in the visual systems of the targeted species, as light penetration in aquatic habitats changes quickly by depth. Five candidate opsin genes were chosen for the analysis: *long-wave sensitive* (*LWS*), *short-wave sensitive 1* (*SWS1*), *short-wave sensitive 2* (*SWS2*), *rhodopsin 2* (*RH2*), and *rhodopsin* (*RH1*) (Table S2).

Using Primer3 software (Koressaar and Remm 2007, Untergasser et al. 2012, Koressaar et al. 2018), primer sets were designed to amplify each of the five genes of interest. The source sequence for *RH1* was identified from a *blastn* (Altschul et al. 1990) search of contigs of unknown identity from a de novo assembly of the *Coregonus artedi* transcriptome (NCBI Bioproject XXXXX) against the nt/nr database (O’Leary et al. 2016). Source sequences for the remaining four genes (*SWS1*, *SWS2*, *LWS*, and *RH2*) were also identified from the *C. artedi* transcriptome, using a *blastn* search of opsin sequences from closely related fishes available from Genbank (Table S5) against the whole transcriptome. After identifying putative opsin sequences from the *C. artedi* transcriptome, these sequences were then searched via *blastn* against the nt/nr database to confirm identity. Primers were selected based on the length of the targeted amplicon, with the goal of amplifying the largest possible fragment from each gene. Following selection, a 22-nucleotide universal tailing sequence was added to the 5’ end of each primer, for compatibility with the Oxford Nanopore Technologies PCR barcoding kit. Primer sequences and annealing temperatures are documented in Table S1.

*DNA Extraction and PCR*

DNA was extracted from gill and/or right pectoral fin tissue using the PureLink Genomic DNA Mini Kit (Invitrogen) following the manufacturer’s protocol. For some samples, lysis was performed following the PureLink Genomic DNA Mini Kit protocol, and lysates were later cleaned using magnetic beads of the HighPrep PCR Clean-up System (MagBio Genomics). Following isolation, genomic DNA was resuspended in 40 μL of either PureLink Genomic Elution Buffer or sterile H_2_O, and stored at -20°C until PCR amplification was performed.

For all samples, each of the five candidate genes was PCR amplified in a separate reaction. Genes were amplified in 25μL reactions containing 2.5 U Accuris Taq DNA Polymerase, 1X Accuris Taq Reaction Buffer, 1 mM dNTPs, 0.4 μM each forward and reverse primers, and ~100 ng genomic DNA. Amplification conditions followed the same general protocol for each gene: an initial denaturation step at 95°C for 3 minutes, followed by 31 rounds of 95°C for 30 seconds, 30 seconds at the specified annealing temperature (Table S2), and a 3 minute extension at 72°C. A final extension at 72°C was carried out for 8 minutes, and resulting amplicons were stored at 4°C. Genes that proved difficult to amplify using the aforementioned protocol were amplified in a 25 μL reaction as follows: 1X LongAmp^®^ *Taq* Master Mix (New England BioLabs), 0.4 μM each of both forward and reverse primers, and ~100 ng genomic DNA. Reaction conditions were maintained the same as above, with the exception of the extension temperature, which was changed from 72°C to 65°C. Following amplification, PCR products were visualized under UV light on a 1% agarose gel stained with 9 μL SYBR^TM^ Safe DNA Gel Stain (Invitrogen).

*Sample-Specific Barcoding*

Following PCR amplification of the five genes of interest, all amplicons from a single individual were pooled into one reaction. Pooled amplicons were cleaned using MagBio magnetic beads, following the manufacturer’s instructions. Post cleanup, total yield per individual was quantified using the dsDNA BR Assay Kit (Invitrogen) on the Qubit 4.0 Fluorometer. Amplicons for each individual were then barcoded using the Oxford Nanopore PCR Barcoding Expansion 1-12. Barcoding was done in a 25 μL reaction as follows: ~15 ng pooled amplicons, 0.2 μM (0.5 μL) Oxford Nanopore Barcode (one of numbers 1-12), and 1X LongAmp^®^ *Taq* Master Mix (New England BioLabs). PCR conditions were as follows: 95°C for three minutes, 15 cycles of 95°C for 15 seconds, 62°C for 15 seconds, and 65°C for 3 minutes, followed by a final extension step of 65°C for 8 minutes. Barcoded amplicons were then purified and quantified as described above using MagBio beads.

*Library Prep and Oxford Nanopore Sequencing*

Following barcoding, amplicons from 12 individuals (i.e., each barcoded with a unique Oxford Nanopore Barcode 1-12) were pooled together in equal ratios to yield 250 ng in 12 μL. Library prep for sequencing was completed using a modified version of the published Oxford Nanopore Protocol for Amplicons by Ligation, using the Ligation Sequencing Kit (SQK-LSK109). DNA repair and end-prep were done in a 15 μL reaction consisting of the 250 ng barcoded amplicons, 0.9 μL NEBNext FFPE DNA Repair Buffer, 0.5 μL NEBNext FFPE DNA Repair Mix, 0.9 μL Ultra II End-prep reaction buffer, and 0.75 μL Ultra II End-prep enzyme mix. The reaction was incubated in a thermocycler for 5 minutes at 65°C followed by 5 minutes at 20°C, after which the sample was again cleaned up using MagBio magnetic beads according to the manufacturer’s protocol, eluting into 16 μL of sterile H_2_O. Adapter ligation was performed using 15 μL of the DNA sample from the previous step, and adding 6.25 μL ligation buffer (Oxford Nanopore Technologies), 2.5 μL NEBNext Quick T4 DNA Ligase, 1.25 μL Adapter Mix (Oxford Nanopore Technologies). The reaction was incubated at room temperature for 10 minutes, and the final product was cleaned up using the following modification of the MagBio beads protocol: 10 μL of resuspended MagBio beads were added to the 25 μL product from the previous step. The solution was mixed well by pipetting, and then incubated on a rotator mixer for 5 minutes to encourage binding of the magnetic beads to the DNA. After 5 minutes, the beads were pelleted on a magnet, and the supernatant was removed. The pellet was washed twice with 200 μL of short fragment buffer (Oxford Nanopore Technologies). Following the second wash, the pellet was dried for ~30 seconds, and the sample was removed from the magnet and resuspended in 15 μL of elution buffer (Oxford Nanopore Technologies). The sample was incubated for 10 minutes on a rotator mixer, following which the beads were once again pelleted on a magnet. 15 μL of eluate containing the fully prepped DNA were removed from the tube. To load the Flongle flow cell, 7.5 μL of sequencing buffer (Oxford Nanopore Technologies), 5 μL of freshly mixed loading beads (Oxford Nanopore Technologies), and 2.5 μL of the prepped DNA were mixed in a new 1.5 mL Eppendorf tube. The flow cell was primed through the sample port with a priming mix consisting of 117 μL of priming buffer and 3 μL of flush tether. Following priming, the freshly mixed solution of sequencing buffer, loading beads, and DNA library was added in a dropwise fashion to the sample port of the flow cell, and a new sequencing run was started. If pore occupancy was initially low (i.e. < 10 pores sequencing), a second library (again consisting of 7.5 μL sequencing buffer, 5 μL loading beads, and 2.5 μL DNA) was added to increase sequencing yield.

In some instances, resequencing of specific genes from certain samples was required, due to low coverage generated on the initial run. In these cases, library prep was completed following the published Oxford Nanopore Protocol for Amplicons by Ligation, using the Ligation Sequencing Kit (SQK-LSK109), yet unlike the protocol detailed above, all reactions were done at the full volume specified (i.e., four times the volumes specified above). The prepared DNA library was then run on a used MinION flow cell. Prior to loading, the used flow cell (FLO-MIN106) was washed using the Oxford Nanopore Flow Cell Wash Kit (EXP-WSH003), and after washing, the flow cell was primed and the library containing barcoded amplicons was loaded, following the manufacturer’s protocol.

In total, seven Flongle flow cells were used for the initial sequencing of amplicons, and three Flongle flow cells and two washed MinION flow cells were used for resequencing.

*Sanger Sequencing*

Sanger sequencing data was generated for *rhodopsin* for 14 of the samples used in the above analyses (3 *Coregonus artedi*, 3 *C. hoyi*, 4 *C. kiyi*, and 4 *C. zenithicus*). *RH1* amplicons were generated in house using the forward and reverse primers 5’- GCATACTCACTCATGGCTGC-3’ and 5’- CTCTGATCCCTGGTTGCTGA - 3’, respectively, and then sent for PCR cleanup and Sanger sequencing to GENEWIZ (South Plainfield, NJ). Reads were generated in both the forward and reverse directions. Low-quality ends of reads were trimmed manually in Geneious Prime v2020.1.2, and base pair calls were visually examined and assigned genotypes. Reads for one individual each of *C. kiyi* were of low quality throughout, and hence were excluded from further analyses. Forward and reverse reads for each individual were aligned in Geneious Prime, and a consensus sequence was generated based on the alignment.

*Data Analysis*

Nanopore data was generated using MinKNOW v3.4.8, and reads were basecalled in real time using Guppy v3.0.6, which is integrated in the MinKNOW platform. For each sequencing run, the high-quality raw fastq files in the fastq_pass folder were converted into a single fasta file. Following this, the script countFasta.pl (<https://github.com/Tininq/countFasta.pl>) was applied to the fasta file, which gave estimates of the total number of base pairs sequenced in each run, as well as the total number of reads and the sequence N50. The average values of each of these metrics were calculated for all Flongle runs.

A representative read length histogram depicting the approximate lengths of reads generated from a single Flongle sequencing run was created from one of the fasta files that was created as detailed above. The python script get_read_lengths.py (<https://github.com/jessiepelosi/FASTA-Scripts/blob/master/get_read_lengths.py>) was used to calculate the length (in bp) of each read in the fasta file, generating a text file containing columns specifying read length and a read ID. This file was manipulated in R using the package ggplot2 v3.3.0 to generate the read length histogram.

An annotated bash script detailing the entire bioinformatic pipeline from read demultiplexing to *F_ST_* calculation is available at: <https://github.com/KrabbenhoftLab/rhodopsin/>. Raw fastq files were demultiplexed and barcodes were trimmed using *guppy_barcoder* v3.2.4 (Oxford Nanopore Technologies), with the options --trim_barcodes and --barcode_kits EXP-PBC001. Nanopore reads from each individual were mapped to version 1 of the *Coregonus* sp. ‘balchen’ genome assembly (De-Kayne et al. 2020; European Nucleotide Archive accession number: GCA_902810595.1) using BWA v0.7.17, with the command *bwa mem* (default parameters; Li 2013). Coordinates of mapped genes are available in Table S6. SAMtools v1.9 was used to convert mapped read files from SAM to BAM format using *samtools view*, and reads were sorted and indexed using *samtools sort* and *samtools index*, respectively (Li et al. 2009), with all default parameters. Mapped reads were visualized in Tablet v1.19.09.03 (Milne et al. 2013), which allowed for a straightforward identification of the genomic locations of the five genes of interest. SNPs were called using BCFtools v1.9 (Li 2010, Li 2011, Danecek et al. 2014), with *bcftools mpileup* (invoking the options -d 1000000 to indicate maximum per-site depth and -R to specify a file containing our regions of interest as determined from visualization in Tablet) and *bcftools call* (invoking the -m option for the multiallelic caller).

Per-site coverage in all individuals was calculated using BEDTools v2.26.0 (Quinlan and Hall 2010), with the command *bedtools coverage* (options: -d). Individual files with per-site coverage were exported into a single concatenated table, with columns specifying gene, sample ID, species ID, and coverage, with each row indicating a distinct nucleotide site. This table was then manipulated in R using the package dplyr v0.8.5, to determine average coverage per gene and average coverage per species.

Consensus sequences of *RH1* generated from both Sanger and nanopore sequencing were compared in Excel to determine the accuracy of nanopore genotyping. At a maximum depth set to 1,000,000x for nanopore data (i.e. maximum depth, as no samples had coverage greater than 32,000x), there was 100% concordance between nanopore and Sanger data, with the exception of a single site that was ambiguous in the Sanger data for one individual. We then performed a rarefaction analysis in which we decreased the maximum per-site depth in a stepwise fashion, using the -d option in *bcftools mpileup* to call SNPs at depths of 2,000, 1,000, 500, 250, 100, 75, 50, and 25x, to determine at which depth nanopore data becomes unreliable for genotyping purposes. *F_ST_* analyses were performed using vcftools v0.1.17 (Danecek et al. 2011) with the option --weir-fst-pop was used to specify the individuals in each population to be analyzed. *F_ST_* analyses were carried out across all four species, as well as in pairwise comparisons between each of the four species.

*Eye diameter measurements and data analysis*

Photos of 32 of the fish used for our analysis (8 *C. artedi*, 8 *C. hoyi*, 9 *C.* kiyi, and 7 *C. zenithicus*) were taken shortly after sampling. Digital landmarks were placed marking eye diameter and head length using the programs TPSDig2 and TPSUtil32 (<https://life.bio.sunysb.edu/morph/soft-dataacq.html>). The ratio of eye diameter to head length was taken for each fish, and plotted in a boxplot by species in R using the package ggplot2 v3.3.0. We analyzed whether there were species-specific differences in eye diameter using a one-way ANOVA, and pairwise comparisons between species were done using Tukey’s honest significance test in R v3.6.3. We also plotted genotype by eye diameter, again using the R package ggplot2 v3.3.0, and performed a logistic regression on these data using the R package stats v3.6.3. The Hosmer and Lemeshow R^2^ value for this logistic regression was also calculated in R.

**Nanopore amplicon library prep and sequencing protocol**

Library Preparation:

1) PCR amplify genes of interest in a 25 μL reaction. Run 5 μL of PCR product on a 1% agarose gel to verify amplicon size and confirm that amplification was successful. Please note – in order to successfully utilize the Oxford Nanopore PCR barcoding expansion kit (step 5), the primers used in this initial PCR step should have the universal Oxford Nanopore primer tail, as shown below:

5’-TTTCTGTTGGTGCTGATATTGC-[project-specific-forward-primer]-3’

5’-ACTTGCCTGTCGCTCTATCTTC-[project-specific-reverse-primer]-3’

1b) Alternatively, if the genes of interest can be amplified simultaneously in a single multiplexed reaction, we recommend this approach, particularly if more than five genes per individual are to be sequenced simultaneously. Skip to step 3.

2) For each individual to be genotyped, pool 10 μL of each amplified gene into a single 1.5 mL Eppendorf tube. (We had five genes per individual, so we ended up with a pool of 50 μL.)

3) Purify the pooled amplicons for each individual. We used magnetic beads as follows:

Pooled amplicons	50 μL

HighPrep^TM^ PCR Clean-up System (MagBio AC-60050)	90 μL

Combine amplicons and magnetic bead solution at a ratio of 1:1.8. Mix by pipetting up and down. Allow beads to bind to amplicons for 5 minutes. Then, pellet on a magnet for ~5 minutes, or until solution is clear and there is a visible pellet. Pipet off supernatant, and add 200 μL of 80% ethanol to wash the pellet. Allow to stand for 30 seconds, then pipet off ethanol. Repeat the ethanol wash once more. After removing the ethanol for a second time, allow the pelleted beads to air dry for 1-2 minutes, but not to the point of cracking. Remove the tube from the magnet, and elute into 15 μL sterile H_2_O. Mix by pipetting up and down, and allow to stand for 5-10 minutes. Pellet on a magnet, and transfer the supernatant into a new 1.5 mL Eppendorf tube – this contains your purified DNA.

4) Quantify the concentration of your purified amplicons. We used a Qubit 4 Fluorometer.

5) Prepare a barcoding PCR – every individual that is to be genotyped in the same sequencing run requires a different barcode. We sequenced 12 at a time, using the Oxford Nanopore Technologies PCR Barcoding Expansion 1-12.

Purified amplicons	50 fmol* in 12 μL

Individual PCR barcode (Oxford Nanopore Technologies EXP-PBC001)	0.5 μL

LongAmp^®^ *Taq* 2X Master Mix (NEB M0287S)	12.5 μL

*While we recommend using 50 fmol of purified PCR product, we note that this can be difficult to calculate if you have amplicons of different lengths. If this is the case, we recommend taking the average length of your amplicons. The following formula can be used to calculate the mass of DNA (in nanograms) to achieve 50 fmol (modified from <https://nebiocalculator.neb.com/#!/dsdnaamt>):

mass (ng) = 50 fmol * 10^-6^ * ((length of amplicons (bp) * 617.96 g/mol) + 36.04 g/mol)

6) Perform the barcoding PCR using the following thermocycler conditions:

| Step Number | Temperature | Time |
| --- | --- | --- |
| 1 | 95°C | 3 minutes |
| 2 | 95°C | 15 seconds |
| 3 | 62°C | 15 seconds |
| 4 | 65°C | 3 minutes** |
| 5 | Go to step 2 15X | -- |
| 6 | 65°C | 8 minutes^+^ |
| 7 | 10°C | Infinite |

**This time is based on the length of the longest amplicon of interest. Typically, we utilize ~1 minute per kb of DNA in the longest amplicon. Our longest amplicon was ~2.5 kb; we chose a 3 minute extension time to ensure the polymerase had adequate time to extend all amplicons.

^+^The time of the final extension is dependent on the length of the amplicons. It can range from 5-10 minutes – we chose 8 minutes, again to ensure the polymerase had adequate time to extend all remaining amplicons.

7) Purify 15 μL of each sample with 27 μL of magnetic beads following the instructions as outlined in step 3. Elute in 10 μL of sterile H_2_O.

8) Quantify the concentration of purified amplicons.

9) Multiplex the barcoded samples that are to be sequenced together into a single 1.5 mL Eppendorf tube. The ideal final concentration of the multiplexed sample should be 250 ng of DNA in 12 μL of sterile H_2_O. (We combined 20.8 ng of each of our 12 individually barcoded samples into this reaction.)

10) Perform DNA repair and end-prep as follows in a sterile 0.2 mL PCR tube:

DNA	250 ng in 12 μL

NEBNext^®^ FFPE DNA Repair Buffer (NEB M6630S)	0.9 μL

NEBNext^®^ FFPE DNA Repair Mix (NEB M6630S)	0.5 μL

NEBNext^®^ Ultra II^TM^ End-prep Reaction Buffer (NEB E7546S)	0.9 μL

NEBNext^®^ Ultra II^TM^ End-prep Enzyme Mix (NEB E7546S)	0.75 μL

Mix by flicking the tube, spin down on a microcentrifuge, and then incubate the reaction in a thermocycler for 5 minutes at 20°C and 5 minutes at 65°C.

11) Purify the reaction using magnetic beads. In this case, use 15 μL of magnetic beads and 15 μL of the end-prepped DNA. In all other regards, follow the steps as outlined in step 3. Elute into 15 μL of sterile H_2_O.

12) Perform adapter ligation as follows in a sterile 1.5 mL Eppendorf tube:

End-prepped DNA from previous step	15 μL

Ligation Buffer (Oxford Nanopore Technologies SQK-LSK109)	6.25 μL

NEBNext^®^ Quick T4 DNA Ligase (E6056S)	2.5 μL

Adapter Mix (Oxford Nanopore Technologies SQK-LSK109)	1.25 μL

Mix by flicking the tube, spin down on a microcentrifuge, and then incubate the reaction at room temperature for 10 minutes.

13) Purify the reaction using magnetic beads. In this case, add only 10 μL of magnetic beads to the 25 μL sample. Mix by pipetting up and down, and incubate the reaction for 5-10 minutes on a rotator mixer. Pellet on a magnet for 5-10 minutes, or until the solution looks clear. Remove the supernatant, and wash the pellet twice using 250 μL of Short Fragment Buffer (if amplicons are larger than 3kb, wash with Long Fragment Buffer to prevent low yield). After removing the buffer for the second time, allow the pellet to air dry for no more than one minute. Remove the tube from the magnetic rack, and elute into 15 μL of Elution Buffer. Pipet up and down to thoroughly mix the sample. Incubate on a rotator mixer for 10 minutes. Re-pellet the beads on a magnet. Transfer 15 μL of the supernatant to a new 1.5 mL Eppendorf tube. This contains your prepped DNA.

Priming and Loading the Flongle Flow Cell:

1) Ensure the Flongle flow cell is properly inserted into your sequencing device. While Oxford Nanopore Technologies recommends running a flow cell check immediately prior to sequencing, we do not recommend this action, as it has resulted in excessive pore loss leading to low sequence yield.

2) Prepare the flow cell priming mix in a 1.5 mL Eppendorf tube as follows:

Flush Buffer (Oxford Nanopore Technologies EXP-FLP002)	117 μL

Flush Tether (Oxford Nanopore Technologies EXP-FLP002)	3 μL

Mix by pipetting up and down.

3) Prime the Flongle flow cell using 120 μL of the priming mix. Draw the priming mix up into a P200 pipette set to 120 μL. Insert the pipet tip directly into the Flongle flow cell sample port. Ensure there are no air bubbles in the pipet tip, and that the tip is flush with the sample port. Slowly dispense all 120 μL into the flow cell, but to avoid the introduction of air bubbles, do not push the pipette past the first stop.

4) Allow the priming mix to sit on the flow cell for 5-10 minutes. While this is occurring, prepare the sample for sequencing.

5) Prepare the library for loading as follows. Ensure the Loading Beads have been mixed thoroughly by pipetting up and down directly before their addition to the sample.

Sequencing Buffer (Oxford Nanopore Technologies SQK-LSK109)	15 μL

Loading Beads (Oxford Nanopore Technologies SQK-LSK109)	10 μL

DNA sample from step 13 of library preparation	5 μL

Mix thoroughly by pipetting up and down prior to loading.

6) Draw up all 30 μL of the prepared library into a P200 pipette. Hover the pipette above the Flongle sample port. In this case, unlike priming, the pipette tip should not be flush with the sample port. Instead, slowly dispense the library from the pipette, allowing it to flow dropwise into the sample port. The liquid should be pulled in to the port, and should not need to be forced in (unlike the priming mix).

7) Once the Flongle flow cell has been loaded, start a sequencing run. We recommend setting the run time for 24 hours, so as to maximize yield.

**Supplemental Tables:**

**Table S1.** Samples used in this study and the geographic location of their collection. Start and end depths indicate the approximate depths (in meters) at which trawls were deployed for each collection.

| **Species** | **Number of individuals** | **Location (latitude, longitude)** | **Start depth (m)** | **End depth (m)** |
| --- | --- | --- | --- | --- |
| *C. artedi* | 2 | 46.80628333, -90.782383333 | 49.9 | 50 |
| *C. artedi* | 3 | 46.95410833, -90.47148333 | 110 | 100 |
| *C. artedi* | 2 | 48.46105833, -88.89977500 | 79.7 | 57.7 |
| *C. artedi* | 1 | 48.43200833, -89.01235000 | 44.5 | 48.2 |
| *C. artedi* | 1 | 48.60125, -88.49655 | 17.2 | 19.7 |
| *C. artedi* | 9 | 48.07695, -89.41105 | 16.5 | 59.2 |
| *C. hoyi* | 2 | 46.89205417, -90.53255833 | 112.5 | 113.5 |
| *C. hoyi* | 5 | 46.88540833, -91.21529167 | 12 | 73.3 |
| *C. hoyi* | 1 | 46.97675833, -90.45365833 | 16.3 | 85 |
| *C. hoyi* | 1 | 46.9022, -90.56715 | 17.1 | 97.3 |
| *C. hoyi* | 10 | 48.07695, -89.41105 | 16.5 | 59.2 |
| *C. kiyi* | 1 | 46.89205417, -90.53255833 | 112.5 | 113.5 |
| *C. kiyi* | 2 | 47.49667500, -89.99912500 | 154 | 154 |
| *C. kiyi* | 2 | 47.15711667, -89.96870833 | 173 | 170 |
| *C. kiyi* | 1 | 47.41659167, -88.46418333 | 195 | 207 |
| *C. kiyi* | 2 | 47.53425000, -90.54291667 | 184 | 185 |
| *C. kiyi* | 4 | 47.1532, -89.97495 | 173 | 169 |
| *C. kiyi* | 8 | 47.0706, -87.1657 | 186 | 183 |
| *C. kiyi* | 1 | 48.46105833, -88.89977500 | 79.7 | 57.7 |
| *C. zenithicus* | 2 | 46.81871667, -91.41867500 | 22.4 | 55.2 |
| *C. zenithicus* | 2 | 47.03325833, -90.60138333 | 42.1 | 48 |
| *C. zenithicus* | 1 | 46.96980833, -90.69405833 | 33.8 | 50.7 |
| *C. zenithicus* | 1 | 46.88540833, -91.2159167 | 12 | 73.3 |
| *C. zenithicus* | 1 | 47.03245000, -90.60183333 | 41.5 | 48.7 |
| *C. zenithicus* | 1 | 46.97315, -91.01 | 17.8 | 50.4 |
| *C. zenithicus* | 8 | 46.88545, -91.21575 | 12 | 73.5 |

**Table S2.** Primers and annealing temperature of PCR amplification (T­_a_), and approximate wavelength of maximum absorption of each opsin analyzed (from Yokoyama et al. 2000).

| **Gene Name** | **Primer Sequences (5’ – 3’)** | **T_a_ (°C)** | **λ_max_ (nm)** |
| --- | --- | --- | --- |
| *Rhodopsin (RH1)* | F: TTTCTGTTGGTGCTGATATTGCGCATACTCACTCATGGCTGC | 57 | 500 |
|  | R: ACTTGCCTGTCGCTCTATCTTCCTCTGATCCCTGGTTGCTGA |  |  |
| *Rhodopsin 2 (RH2)* | F: TTTCTGTTGGTGCTGATATTGCGTTCAGAGCCAGATCAGAT | 56 | 470-510 |
|  | R: ACTTGCCTGTCGCTCTATCTTCATCTCGTGATCCAATCTCAA |  |  |
| *Long-wave sensitive (LWS)* | F: TTTCTGTTGGTGCTGATATTGCTCTTCTGCTACATTTTCGTG | 55 | 510-560 |
|  | R: ACTTGCCTGTCGCTCTATCTTCTAGGTTTCGTGGGTAATGTT |  |  |
| *Short-wave sensitive 1 (SWS1)* | F: TTTCTGTTGGTGCTGATATTGCAGACCTGAATGTGACTTTTA | 54 | 360-430 |
|  | R: ACTTGCCTGTCGCTCTATCTTCGAAGTCCTTTGCCATCTTG |  |  |
| *Short-wave sensitive 2 (SWS2)* | F: TTTCTGTTGGTGCTGATATTGCAGAAGGAGGTCACCAAGAT | 52 | 440-460 |
|  | R: ACTTGCCTGTCGCTCTATCTTCGCTGCTGAAGTCTAAATTTGA |  |  |

**Table S3.** Mean coverage of all genes by species.

| **Species** | **Mean coverage (±SD; min-max)** |
| --- | --- |
| *C. artedi* | 3368.55x ± 5386.53x; 58.07-22976.74x |
| *C. hoyi* | 4520.00x ± 5531.35x; 46.71-20531.73x |
| *C. kiyi* | 1988.36x ± 2965.37x; 44.06-15786.71x |
| *C. zenithicus* | 2970.53x ± 4704.10x; 10.47-31158.31x |

**Table S4.** Cost analysis of a single Oxford Nanopore Flongle genotyping run. Final cost does not include the one-time purchases of the Flongle Flow Cell Adapter nor the MinION sequencing device, which are required for Flongle sequencing runs.

| **Component** | **Vendor** | **Cost per Flongle run (USD)** |
| --- | --- | --- |
| Ligation sequencing kit (SQK-LSK109) | Oxford Nanopore Technologies | $24.96 |
| PCR barcoding expansion 1-96 (EXP-PBC096) | Oxford Nanopore Technologies | $42.50 |
| HighPrep^TM^ PCR Clean-up System (AC-60500) | MagBio Genomics | $82.82 |
| LongAmp^®^ *Taq* 2X Master Mix (M0287L) | New England BioLabs | $51.55 |
| NEBNext^®^ FFPE DNA Repair Mix (M6630L) | New England BioLabs | $1.50 |
| NEBNext^®^ Ultra^TM^ II End Repair/dA-Tailing Module (E7546L) | New England BioLabs | $2.07 |
| NEBNext^®^ Quick Ligation Module (E6056L) | New England BioLabs | $6.24 |
| Flongle Flow Cell | Oxford Nanopore Technologies | $90.00 |
| **Total Cost:** |  | **$301.64** |

**Table S5.** Sequences used to locate putative opsins in the *C. artedi* transcriptome.

| **Gene** | **Species** | **Genbank accession number** |
| --- | --- | --- |
| *SWS1* | *Oncorhynchus mykiss* | XM_021563440 |
| *SWS2* | *Salvelinus alpinus* | XM_023995084 |
| *LWS* | *Salmo salar* | XM_014133083 |
| *RH2* | *Oncorhynchus tshawytscha* | XM_024427705 |

**Table S6.** Locations of each of the opsin amplicons within version one of the *C. sp ‘balchen’* genome (De-Kayne et al. 2020).

| **Gene** | **Chromosome ID** | **Location (bp)** |
| --- | --- | --- |
| *RH1* | ENA\|LR664373\|LR664373.1 | 15258695-15259457 |
| *LWS* | ENA\|LR664353\|LR664353.1 | 18319277-18321155 |
| *SWS2* | ENA\|LR664353\|LR664353.1 | 19555690-19556649 |
| *SWS1* | ENA\|LR664371\|LR664371.1 | 18549973-18550787 |
| *RH2* | ENA\|LR664355\|LR664355.1 | 39573974-39576052 |

**Supplemental Figures:**

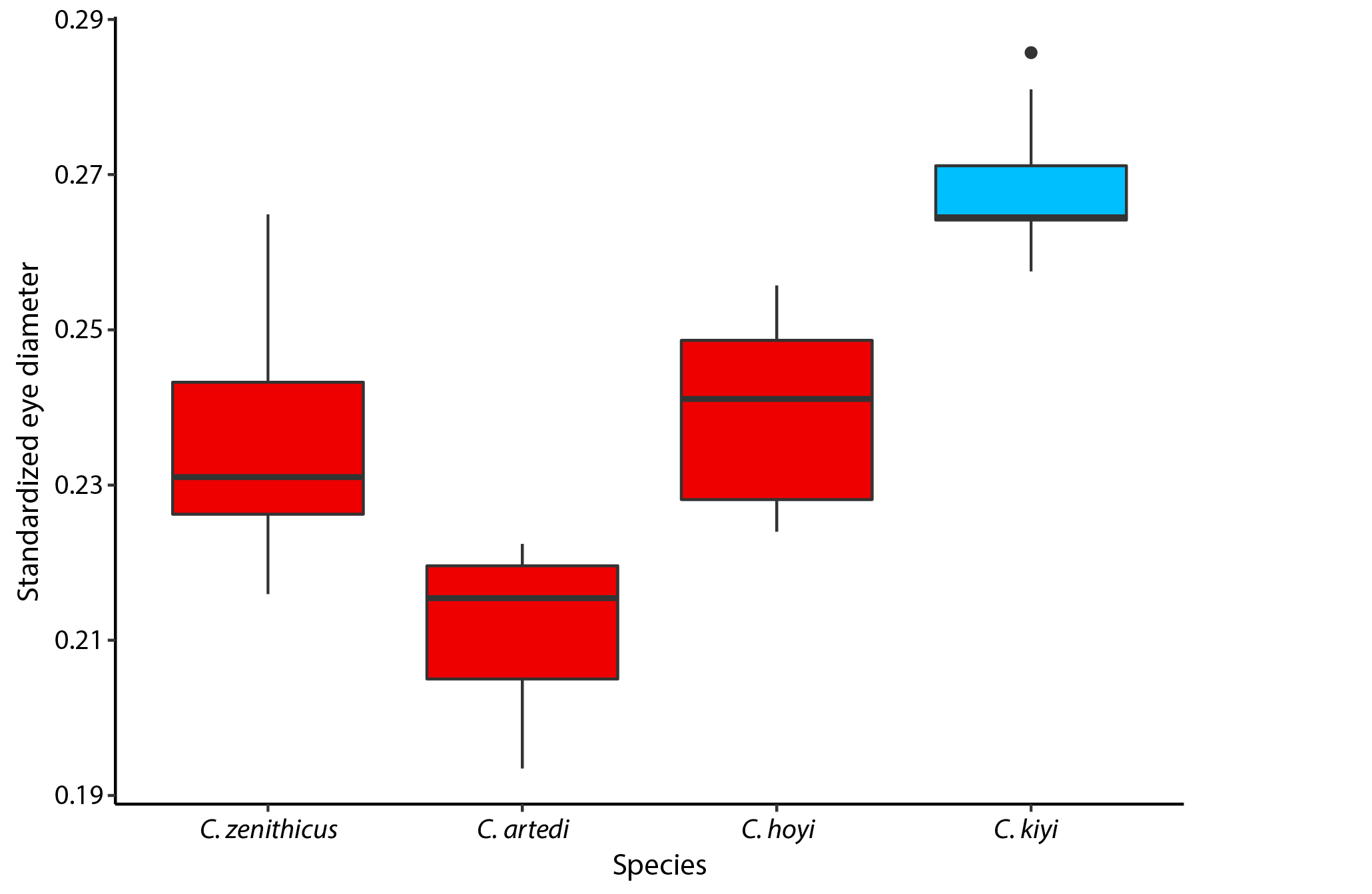

**Figure S1**. Boxplot of eye diameter (standardized by head length) vs. species. Boxes are colored based on the most common amino acid found at *rhodopsin* residue 261, with red representing the red-shifted tyrosine, and blue representing the blue-shifted phenylalanine. Based on Tukey’s HSD test for multiple comparisons of means, significant differences in eye diameter were found between: *C. zenithicus* and *C. artedi* (F_3.28_ = 30.92, p < 0.01), *C. zenithicus* and *C. kiyi* (F_3.28_ = 30.92, p < 0.001), *C. hoyi* and *C. artedi* (F_3.28_ = 30.92, p < 0.001), *C. kiyi* and *C. artedi* (F_3.28_ = 30.92, p < 0.001), and *C. kiyi* and *C. hoyi* (F_3.28_ = 30.92, p < 0.001).

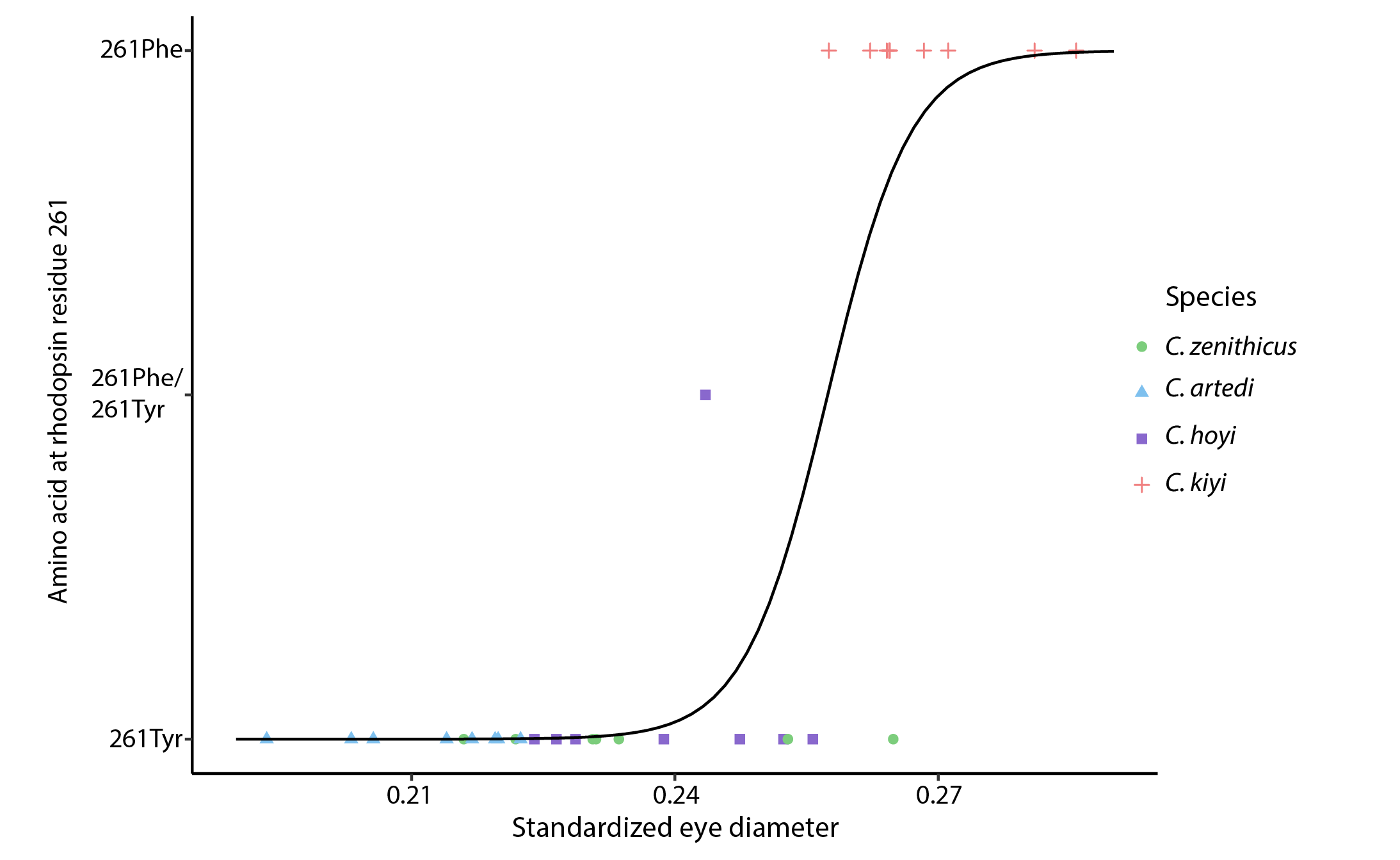

**Figure S2.** Logistic regression plot of amino acid at *rhodopsin* residue 261 vs. eye diameter (standardized by head length). A logistic regression has been applied to the plot; the Hosmer and Lemeshow R^2^ value for this regression was found to be 0.698, indicating that eye diameter accounts for 69.8% of the variability in genotype at this locus.
